# Parallel ZNF598 and GIGYF2 pathways mediate conserved collision-dependent mRNA decay with distinct endonucleolytic features

**DOI:** 10.64898/2026.01.01.697312

**Authors:** Satoshi Hashimoto, Jennifer Sauerland, Jiahuan Zeng, Chisato Kikuguchi, Toshifumi Inada

**Affiliations:** Division of RNA and Gene Regulation, Institute of Medical Science, The University of Tokyo, Minato-Ku, 108-8639, Japan; Gene Center and Department of Biochemistry, University of Munich LMU, Feodor-Lynen-Str. 25, 81377 Munich, Germany

## Abstract

Quality control of aberrant mRNAs is essential for maintaining cellular homeostasis. Ribosome collisions caused by translational stalling trigger mRNA surveillance pathways that promote degradation of faulty transcripts. In yeast, two major collision-responsive pathways have been proposed: the Hel2–Cue2 pathway and the Syh1–Xrn1 pathway, yet their functional relationship and conservation in mammals have remained unclear. Here, we demonstrate that these two pathways act redundantly to degrade collision-inducing mRNAs in both yeast and mammalian cells. In yeast, we dissect the contribution of Cue2-mediated endonucleolytic cleavage and show that eS7 polyubiquitination–dependent cleavage plays a dominant role in Hel2–Cue2 pathway. In mammals, although the proposed Cue2 homolog N4BP2 is dispensable, we provide the first direct evidence of collision-induced endonucleolytic mRNA cleavage. Together, our findings reveal both conserved and divergent features of ribosome collision–coupled mRNA quality control and expand our understanding of mRNA surveillance and potential therapeutic mRNA implications.

## INTRODUCTION

Ribosomes do not always translate mRNAs smoothly. Certain sequence contexts, such as tandem rare codons in yeast(Dimitrova et al., 2009) or poly(A) stretches in yeast and mammalian cells, can induce ribosome stalling during elongation(Dimitrova et al., 2009; Inada & Aiba, 2005; Juszkiewicz & Hegde, 2017; Sundaramoorthy et al., 2017). When elongation is severely impaired, trailing ribosomes collide with a stalled leading ribosome, resulting in the formation of a characteristic disome structure(Ikeuchi et al., 2019; Matsuo et al., 2020; Tesina et al., 2020). Ribosome collisions act as a key signal to activate quality control pathways that target both nascent polypeptides and their associated mRNAs in a collision-dependent manner. Defective nascent polypeptides are eliminated by the ribosome-associated quality control (RQC) pathway(Brandman et al., 2012; Hashimoto et al., 2020; Juszkiewicz & Hegde, 2017; Matsuo et al., 2017; Narita et al., 2022; Sundaramoorthy et al., 2017), whereas aberrant mRNAs are degraded through no-go decay (NGD) (D’Orazio et al., 2019; Ikeuchi et al., 2019; Müller et al., 2025; Tomomatsu et al., 2023).

The molecular mechanisms of NGD have been extensively characterized in yeast. Ribosome collisions induce endonucleolytic cleavage of the mRNA, generating fragments that are subsequently degraded by the 5′–3′ exonuclease Xrn1 or the 3’-5’ exonuclease complex Ski–exosome complex (Doma & Parker, 2006). We previously demonstrated that NGD involves two mechanistically distinct modes of endonucleolytic cleavage: NGD–RQC⁺, in which cleavage occurs within the collided ribosome, and NGD–RQC⁻, in which cleavage occurs upstream of the collision site (Ikeuchi et al., 2019; Tomomatsu et al., 2023). In the NGD–RQC⁺ pathway, ribosome collisions are first sensed by the E3 ubiquitin ligase Hel2, which catalyses K63-linked polyubiquitination of the small ribosomal subunit protein uS10 of the leading ribosome (Matsuo et al., 2017; Saito et al., 2015). The ubiquitinated uS10 is recognized by Cue3/Rqt3 and Rqt4, leading to recruitment of the Slh1/Rqt2 helicase, which dissociates the leading stalled ribosome (Best et al., 2023; Matsuo et al., 2017, 2020, 2023). Following ribosome dissociation, the endonuclease Cue2 cleaves the mRNA within the collided ribosome. When collided ribosomes are not dissociated, an alternative pathway, termed NGD–RQC⁻, is engaged (Tomomatsu et al., 2023). In this pathway, the E3 ligase Not4 first mono-ubiquitinates eS7A at the four lysine residues, which is subsequently extended by Hel2 to form K63-linked polyubiquitin chains (Ikeuchi et al., 2019). Cue2 is then recruited to polyubiquitinated eS7A via its N-terminal ubiquitin-binding CUE-D1 and CUE-D2 domains and cleaves the mRNA upstream of the disome unit (Tomomatsu et al., 2023). Thus, in yeast, Cue2-mediated endonucleolytic cleavage represents a central feature of NGD, operating through two distinct ubiquitin-dependent mechanisms.

In parallel to the Hel2–Cue2 axis, an independent pathway contributes to collision-dependent mRNA decay by promoting translational repression and exonucleolytic degradation. In yeast, colliding ribosomes are proposed to recruit Syh1 and its paralog Smy2, which facilitate decapping and Xrn1-dependent degradation of aberrant mRNAs (Veltri et al., 2022). In mammalian cells, homologous factors have been identified, including ZNF598 as the Hel2 (Hashimoto et al., 2020; Juszkiewicz et al., 2018; Sundaramoorthy et al., 2017) and GIGYF2 as a functional analogue of the Syh1–Smy2 axis (Ash et al., 2010; Hickey et al., 2020; Juszkiewicz et al., 2020; Weber et al., 2020). However, the conservation of NGD mechanisms between yeast and mammals remains incompletely understood. In particular, while N4BP2 has been proposed as a mammalian homolog of the yeast endonuclease Cue2 (Glover et al., 2020; Nicholson-Shaw et al., 2025; Saito et al., 2022), its contribution to collision-dependent mRNA decay has not been clearly established. Moreover, direct evidence for endonucleolytic cleavage of collision-inducing mRNAs in mammalian cells has been lacking.

Here, we demonstrate that the Hel2/ZNF598 and Syh1–Smy2/GIGYF2 pathways act redundantly to promote the decay of collision-inducing mRNAs in both yeast and mammalian cells. In yeast, we further dissect the contribution of Cue2-mediated endonucleolytic cleavage pathways to mRNA decay and show that eS7 polyubiquitination–dependent NGD–RQC⁻ cleavage contributes dominantly to mRNA decay in the absence of the Syh1–Smy2 pathway. In mammalian cells, we find that the proposed Cue2 homolog N4BP2 is dispensable for NGD. Instead we will provide the first direct evidence of collision-induced endonucleolytic mRNA cleavage by SMG6, an essential NMD factor. Overall, these findings reveal both conserved and divergent features of ribosome collision–coupled mRNA quality control.

## RESULTS

### Hel2 and Syh1 act independently to degrade aberrant mRNA in yeast

No-go decay (NGD) involves two mechanistically distinct modes of endonucleolytic cleavage; however, the contribution of those endonucleolytic cleavage pathways in mRNA decay remains unclear. Moreover, the functional relationship between the Hel2–Cue2 and Syh1–Xrn1 pathways in collision-dependent mRNA decay has not been fully addressed. To accurately assess mRNA stability, we measured reporter mRNA half-lives using a *GAL1* promoter–based glucose transcriptional shutoff system (Fig. 1B). In this system, reporter transcription is active in the presence of galactose and is rapidly repressed upon glucose addition (Coller, 2008; Lohr et al., 1995; Yano & Fukasawa, 1997). Using this approach, we measured the half-life of the *GFP-R12-HIS3* reporter, a well-established substrate of no-go decay (NGD) in yeast in which a tandem rare codon sequence efficiently induces ribosome collisions (Dimitrova et al., 2009; D’Orazio et al., 2019; Ikeuchi et al., 2019; Kuroha et al., 2010; Tomomatsu et al., 2023; Veltri et al., 2022). Measurements were performed in strains lacking key components of the two collision-response pathways, namely Hel2 and Syh1–Smy2. Deletion of *HEL2* resulted in a modest but not statistically significant increase in reporter mRNA stability, while deletion of *SYH1-SMY2* had little effect compared with wild-type W303 cells (Fig. 1C, D). These results suggested that the two pathways might function redundantly in degrading collision-inducing mRNAs. Consistent with this interpretation, combined deletion of *HEL2* and *SYH1-SMY2* led to a pronounced stabilization of the *GFP-R12-HIS3* reporter, relative to either single deletion (Fig. 1C, D). Importantly, this effect was specific to aberrant mRNAs, as no stabilisation was observed for the non-stalling *GFP-R0-HIS3* reporter (Fig. 1E, F). These findings are consistent with previous observations using the minCGA reporter containing a CGA_12_ sequence within the ORF (Veltri et al., 2022). Since NGD is tightly coupled to translation, we next examined whether Syh1 associates with translating ribosomes. Polysome profiling revealed that Syh1 was broadly distributed across polysome fractions even in the absence of Hel2 (Fig. 1G), supporting a model in which the Hel2-Cue2 and Syh1-Xrn1 pathways act independently to promote degradation of collision-inducing mRNAs.

**Fig. 1.**
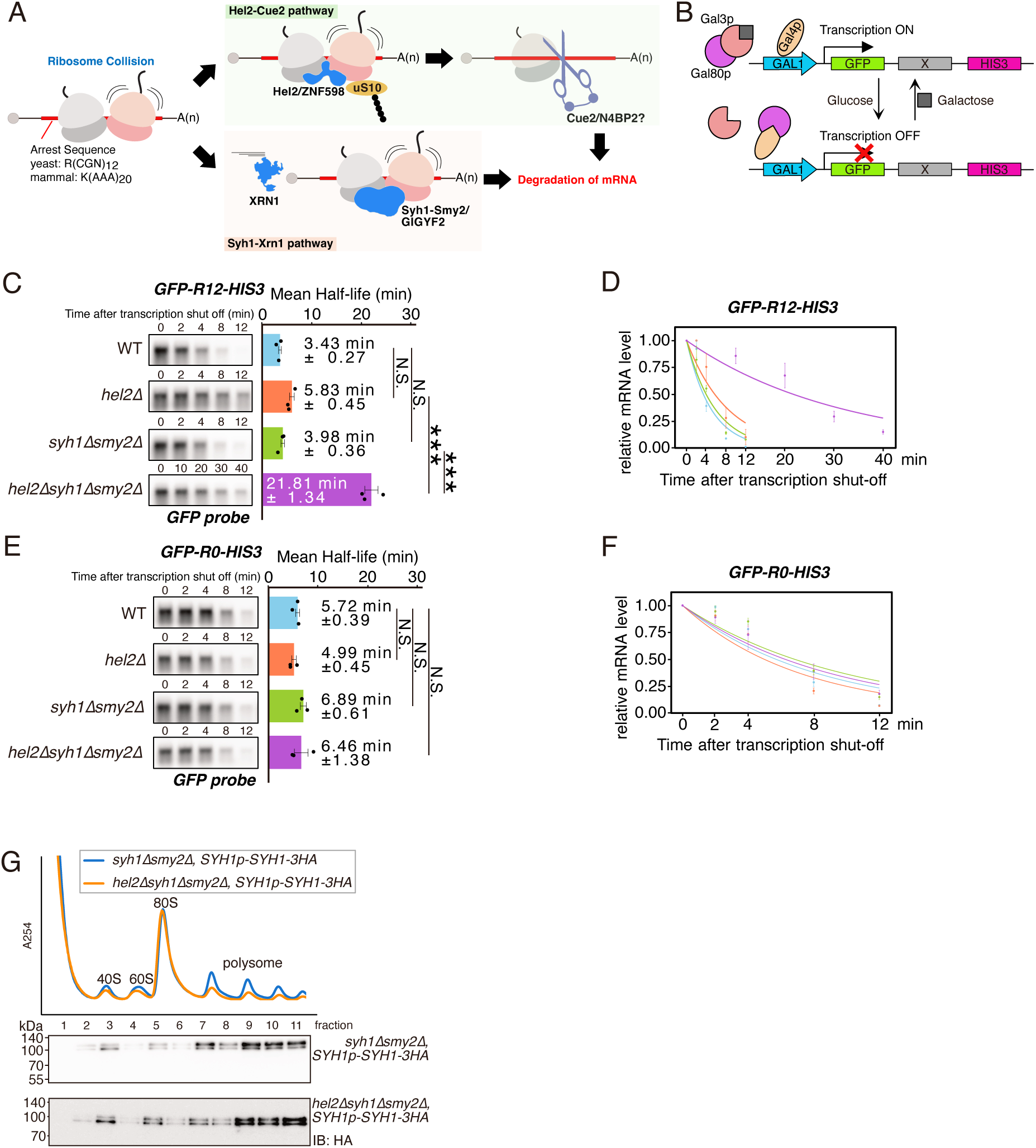
Hel2 and Syh1 act in parallel to degrade aberrant mRNAs in yeast. **A.** Schematic summary of the two putative pathways involved in aberrant mRNA degradation in yeast and mammals. Key regulatory factors of the Hel2–Cue2 and Syh1–Xrn1 pathways are shown (yeast/mammal). **B.** Schematic of the GAL1 promoter–based glucose shut-off system used to measure mRNA half-life in yeast. Reporter transcription is active in the presence of galactose and rapidly repressed upon glucose addition. **C–F.** Half-life measurements of reporter mRNAs in the indicated deletion backgrounds. **C, E (left):** Northern blot analysis of the indicated reporters detected with a GFP probe. **C, E (right):** Estimated mRNA half-lives calculated from relative mRNA levels. Each dot represents a half-life from individual experiment (n = 3); ; bars indicate mean values, and error bars represent S.E. Statistical analysis was performed using the Benjamini–Hochberg method. Adjusted *p* values are indicated as follows: *** *p*adj < 0.001; ** *p*adj < 0.01; * *p*adj < 0.05; N.S., not significant. **D, F.** Quantification of relative mRNA levels from northern blots shown in C and E. Regression lines were generated by nonlinear least-squares fitting to an exponential decay model of normalized RNA abundance over time. Colours correspond to those used in C and E. **G.** Sucrose density gradient–western blot (SDG-WB) analysis of the indicated strains expressing *SYH1–3HA*. Blots were probed with an anti-HA antibody. Representative results from three independent experiments are shown.

### eS7 polyubiquitination-mediated mRNA cleavage by Cue2 promotes mRNA decay independently of Syh1

Endonucleolytic cleavage by Cue2 is a central feature of no-go decay (NGD) in yeast (D’Orazio et al., 2019; Ikeuchi et al., 2019; Kuroha et al., 2010; Tomomatsu et al., 2023; Veltri et al., 2022). To define the contribution of Cue2 to mRNA degradation, we measured reporter mRNA half-lives in strains lacking *CUE2*. Deletion of *CUE2* alone did not significantly alter mRNA stability (Fig. 2A, B), similar to what we observed in *HEL2* deletion strain (Fig. 1C, D). This result is consistent with previous findings that Cue2 functions downstream of Hel2-dependent ribosome ubiquitination (Tomomatsu et al., 2023). In contrast, simultaneous deletion of *CUE2* together with *SYH1-SMY2* resulted in a pronounced increase in the half-life of the aberrant mRNA compared with either single deletion (Fig. 2A, B). This stabilization was specific to aberrant mRNAs and was not observed for non-stalling mRNAs (Fig. 2C, D). Polysome profiling further showed that Cue2 association with ribosomes was unaffected by deletion of *SYH1-SMY2* (Fig. 2E), supporting that Cue2 functions independently of the Syh1-Xrn1 pathway.

**Fig. 2.**
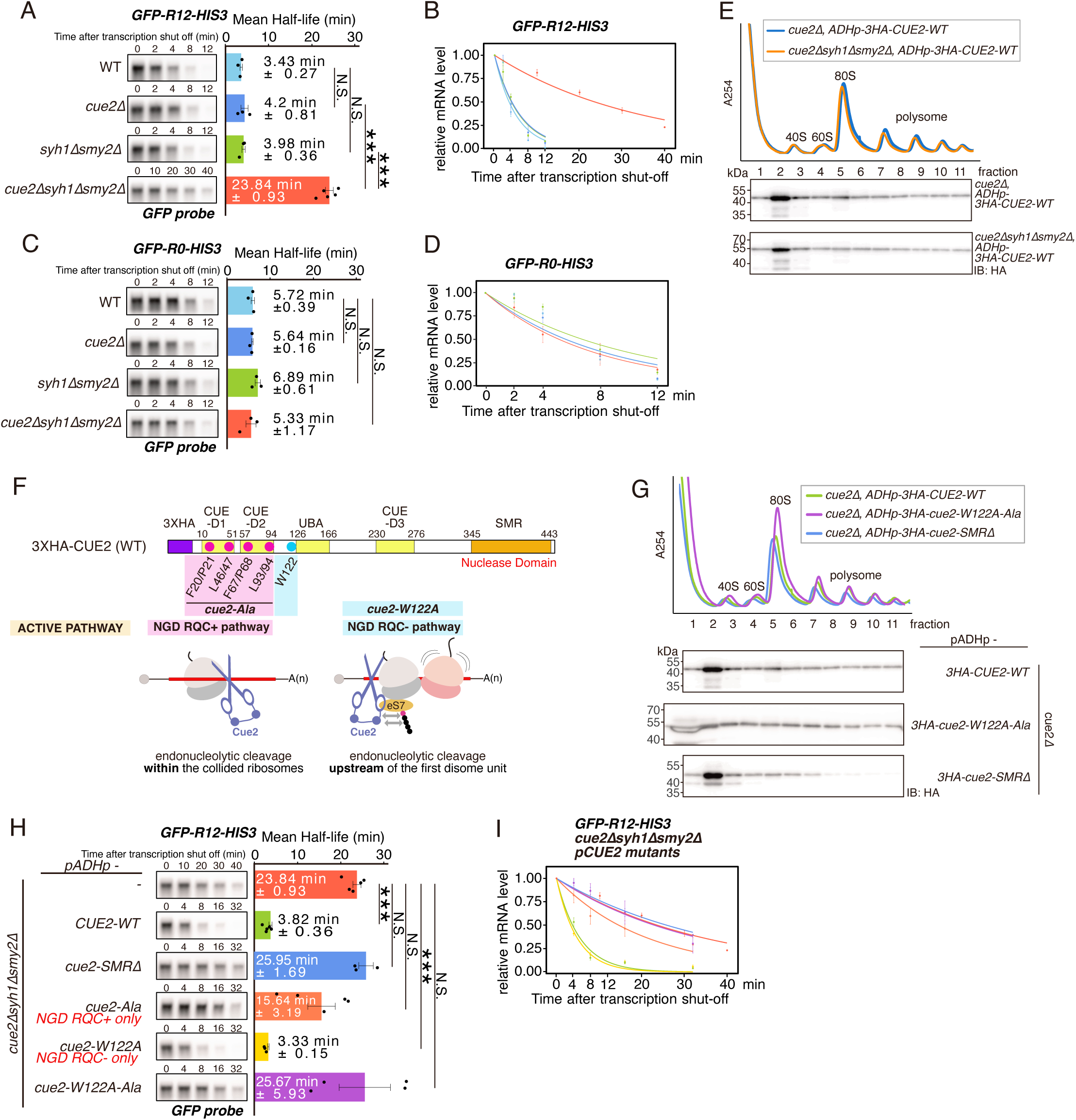
Cue2 and Syh1 act in parallel to degrade aberrant mRNAs in yeast. **A–D, H–I.** Half-life measurements of reporter mRNAs in the indicated deletion backgrounds. **A, C, H (left):** Northern blot analysis of the indicated reporters detected with a GFP probe. **A, C, H (right):** Estimated mRNA half-lives calculated from relative mRNA levels. Each dot represents a half-life from individual experiment (n = 3–5); bars indicate mean values, and error bars represent S.E. Statistical analysis was performed using the Benjamini–Hochberg method. Adjusted *p* values are indicated as follows: *** *p*adj < 0.001; ** *p*adj < 0.01; * *p*adj < 0.05; N.S., not significant. **B, D, I.** Quantification of relative mRNA levels from the northern blots shown in A, C, and H. Regression lines were generated by nonlinear least-squares fitting to an exponential decay model of normalized RNA abundance over time. Colors correspond to those used in A, C, and H. **F.** Schematic illustration of Cue2 domain map and its two modes of endonucleolytic cleavage, NGD-RQC⁺ and NGD-RQC⁻. **E and G.** Sucrose density gradient–western blot (SDG-WB) analysis of the indicated strains expressing Cue2 mutants. Blots were probed with an anti-HA antibody. Representative results from three independent experiments are shown.

Cue2 contains a conserved Small MutS-Related (SMR) nuclease domain which is essential for endonucleolytic cleavage of the aberrant mRNA (Saito et al., 2022; Veltri et al., 2022) (Fig. 2F). To test the functional requirement of this domain, we performed complementation assays in a *cue2Δ syh1Δ smy2Δ* background (Fig. 2H, I). Expression of wild-type Cue2 (CUE2-WT) fully restored normal mRNA decay, whereas an SMR-deleted mutant (cue2–SMRΔ) failed to complement the decay defect (Fig. 2H, I), demonstrating that Cue2 nuclease activity is essential for degradation of aberrant mRNAs. Cue2 mediates two distinct modes of endonucleolytic cleavage: NGD-RQC^+^, in which mRNA is cleaved within collided ribosomes, and NGD-RQC^−^, in which cleavage occurs upstream of collided ribosomes in an eS7 polyubiquitination-dependent manner (Tomomatsu et al., 2023) (Fig2.F). To assess the contribution of these two cleavage modes to mRNA decay, we expressed mode-specific Cue2 mutants. Expression of an NGD-RQC^+^-only mutant (cue2-Ala) partially restored mRNA decay, whereas an NGD-RQC^−^-only mutant (cue2-W122A) fully rescued the decay defect to wild-type levels. As expected, a double-inactive mutant (cue2-W122A-Ala) failed to restore mRNA degradation (Fig. 2H, I). Ribosome association of the double-inactive mutant (cue2-W122A-Ala) was comparable to that of wild-type Cue2, whereas the Cue2-SMRΔ mutant exhibited reduced ribosome binding (Fig. 2G), suggesting that the SMR domain itself contributes to ribosome association. This reduction is consistent with previous observations that MutS, an SMR domain–containing protein, associates with collided ribosomes in *Bacillus subtilis* (Park et al., 2023). Together, these results demonstrate that eS7 polyubiquitination–dependent mRNA cleavage by Cue2 is a major contributor to aberrant mRNA degradation and functions independently of the Syh1-Xrn1 pathway.

### ZNF598 and GIGYF2 act redundantly in mammalian collision-dependent mRNA decay

To investigate whether collision-dependent mRNA decay is conserved in mammalian cells, we employed a Tet-off transcriptional shutoff system to measure reporter mRNA stability (Gossen & Bujard, 1992). In this system, transcription of the reporter gene is driven by a tetracycline-responsive promoter and can be rapidly and specifically repressed by the addition of doxycycline, allowing direct measurement of reporter mRNA decay kinetics without affecting the endogenous transcription level (Fig. 3A). To test whether the degradation mechanisms identified in yeast are conserved in mammals, we generated a series of knockout (KO) cell lines targeting mammalian homologs of key yeast collision-response factors: ZNF598 (Hel2 homolog) (Hashimoto et al., 2020; Juszkiewicz et al., 2018; Sundaramoorthy et al., 2017), N4BP2 (proposed Cue2 homolog) (Glover et al., 2020; Nicholson-Shaw et al., 2025; Saito et al., 2022)., and GIGYF2 (Syh1 homolog) (Ash et al., 2010; Fukao et al., 2021; Hickey et al., 2020; Juszkiewicz et al., 2020; Morita et al., 2012; Veltri et al., 2022; Weber et al., 2020) (Fig. 3B). A doxycycline-responsive reporter was stably integrated into the genome of each KO cell line.

**Fig. 3.**
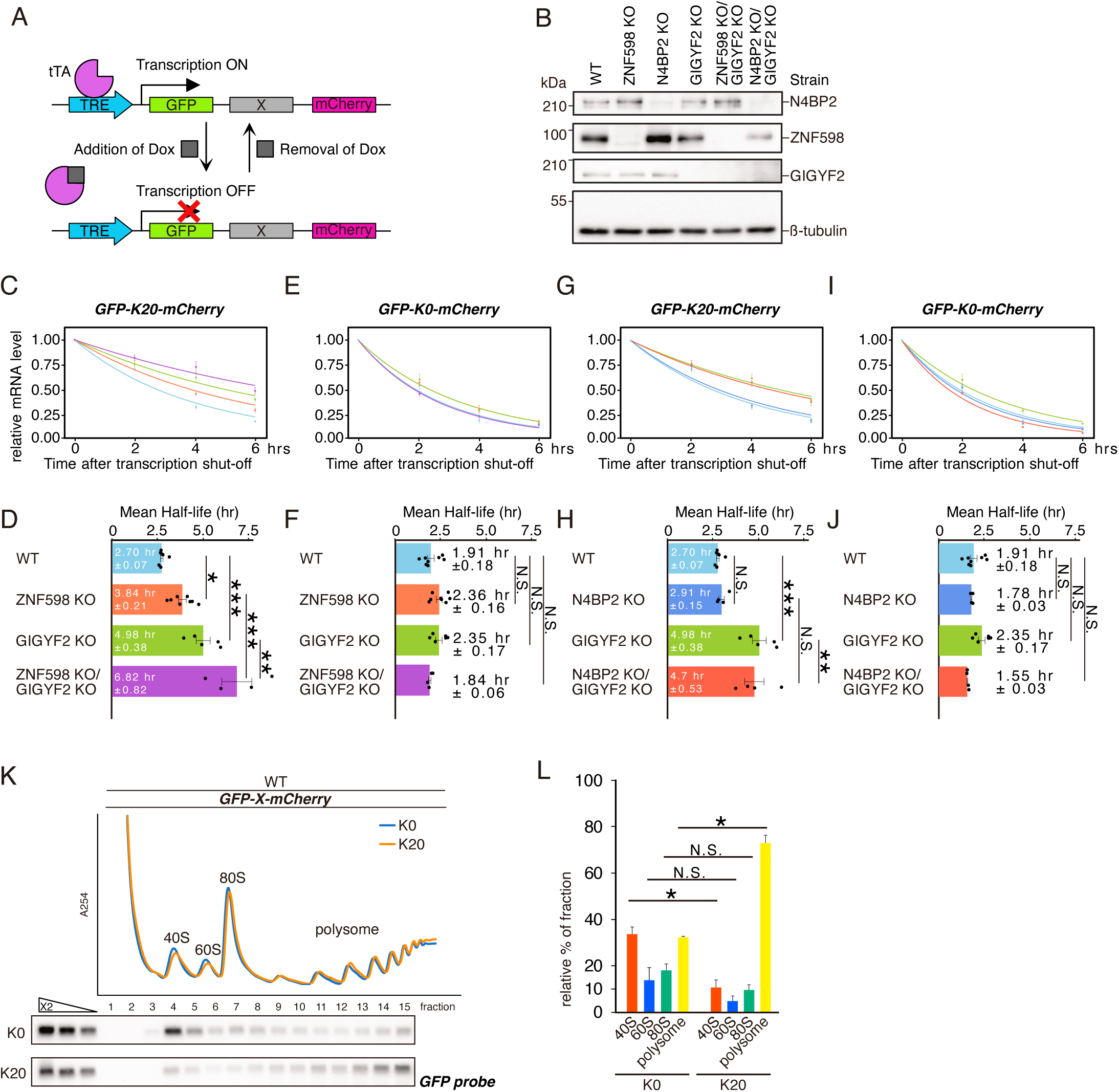
ZNF598 and GIGYF2 act redundantly to degrade aberrant mRNAs in mammalian cells. **A.** Schematic of the Tet-off reporter system. The tetracycline transactivator (tTA) binds to the tetracycline response element (TRE) and activates transcription in the absence of doxycycline (Dox). Upon Dox addition, tTA dissociates from the TRE, resulting in transcriptional shutoff of the reporter mRNA. **B.** Western blot analysis of knockout (KO) cell lines used for mRNA half-life measurements. **C, E, G, I.** Relative mRNA levels of the *GFP-X-mCherry* reporters normalized to the level of 0 h time point. **D, F, H, J.** Estimated mRNA half-lives calculated from the data shown in C, E, G, and I. Each dot represents a half-life from individual experiment (n = 3–8); bars indicate mean values, and error bars represent S.E. Statistical analysis was performed using the Benjamini–Hochberg method. Adjusted *p* values are indicated as follows: *** *p*adj < 0.001; ** *p*adj < 0.01; * *p*adj < 0.05; N.S., not significant. **K.** Sucrose density gradient–northern blot (SDG-NB) analysis of *GFP-X-mCherry* reporters. Distribution of reporter mRNAs across gradient fractions was detected by northern blotting using a GFP probe. Representative results from two independent experiments are shown. **L.** Quantification of reporter mRNA distribution across sucrose gradient fractions shown in K. Statistical significance was assessed using Student’s *t*-test. *p* < 0.05 (*); N.S., not significant.

Using these cell lines, we quantified reporter mRNA half-lives by RT–qPCR following transcriptional shutoff. When a collision-inducing reporter containing a polybasic stall sequence (*GFP-K(AAA)_20_-mCherry*) (Hashimoto et al., 2020; Juszkiewicz et al., 2018; Sundaramoorthy et al., 2017) was expressed, single deletion of either ZNF598 or GIGYF2 resulted in a modest increase in reporter mRNA half-life. In contrast, simultaneous deletion of both ZNF598 and GIGYF2 led to a pronounced stabilization of the aberrant reporter mRNA compared with either single deletion (Fig. 3C, D). This phenotype closely mirrors observations in yeast and supports a model in which two genetically separable pathways act redundantly to promote degradation of collision-inducing mRNAs. Importantly, no significant changes in mRNA stability were observed for the non-stalling control reporter (*GFP-K_0_-mCherry*) in any genetic background (Fig. 3E, F).

We next asked whether N4BP2 functions as a mammalian homolog of the yeast endonuclease Cue2. Deletion of N4BP2 alone did not extend the half-life of the collision-inducing reporter, consistent with observations in yeast where Cue2 deletion alone has little effect on mRNA stability (Fig.2A). However, unlike the yeast Cue2 pathway, simultaneous deletion of N4BP2 and GIGYF2 did not further stabilize the *GFP-K_20_-mCherry* reporter compared with GIGYF2 single knockout (Fig. 3G, H). The half-life of the non-stalling *GFP-K_0_-mCherry* reporter remained unchanged across all knockout backgrounds (Fig. 3I, J). These findings indicate that, although redundant collision-dependent mRNA decay pathways are conserved in mammalian cells, N4BP2 does not function as a Cue2 in this context (Glover et al., 2020; Nicholson-Shaw et al., 2025; Saito et al., 2022).

Interestingly, we observed that the collision-inducing *GFP-K20-mCherry* reporter exhibited a slightly longer half-life than the non-stalling *GFP-K0-mCherry* reporter (Fig3. D and F). This counterintuitive stabilisation can be explained by increased ribosome stacking on the collision-inducing reporter relative to the non-stalling reporter. A higher density of stalled ribosomes on a mRNA has been proposed to protect transcripts from decay by increasing ribosome occupancy and thereby shielding the mRNA from decay factors (Bicknell et al., 2024; Dilrangi et al., 2025; El Fatimy et al., 2016; Ripin & Parker, 2022, 2023; Sheth & Parker, 2003). To test this hypothesis, we analysed ribosome association of the reporter mRNAs using sucrose density gradient fractionation followed by Northern blotting (SDG-NB). Consistent with our hypothesis, the *GFP-K20-mCherry* reporter mRNA was more enriched in polysome fractions than the *GFP-K0-mCherry* reporter (Fig. 3K, L). These results support a model in which increased ribosome occupancy contributes to the stabilization of collision-inducing transcripts.

### Ribosome slippage on the K20 sequence induces SMG6-dependent endonucleolytic cleavage

Although N4BP2 does not appear to function as a Cue2 in mammalian cells, whether collision-dependent endonucleolytic cleavage occurs during mammalian no-go decay (NGD) remains unclear. To date, endonucleolytic intermediates of mammalian NGD have not been clearly documented. Previous studies of mammalian NGD have primarily relied on non-stop decay (NSD)-type reporters and were not performed in an XRN1-deficient background, conditions that may have precluded detection of endonucleolytically cleaved intermediates (Glover et al., 2020; Ishibashi et al., 2024; Nicholson-Shaw et al., 2025). To address this possibility, we established an experimental system designed to capture endonucleolytic cleavage intermediates.

To directly detect potential cleavage intermediates, we generated XRN1 knockout cells to inhibit 5′–3′ exonucleolytic decay, thereby stabilizing 3′ RNA intermediates generated by endonucleolytic cleavage. In this background, we expressed *GFP-K0-mCherry* or *GFP-K20-mCherry* reporters and detected reporter mRNAs using probes targeting either the 5′ region (GFP probe) or the 3′ region (mCherry probe) (Fig. 4A). When probed with the GFP probe, only full-length mRNA was detected for both reporters (Fig. 4A, lanes 3 and 4). In striking contrast, probing for mCherry revealed a distinct 3′ NGD intermediate specifically in cells expressing the K20 reporter (Fig. 4A, lane 8). Notably, this intermediate was not detected for the non-stalling K0 reporter (Fig. 4A, lane 7). To our knowledge, this constitutes the first direct detection of a 3′ NGD intermediate in mammalian cells.

**Fig. 4.**
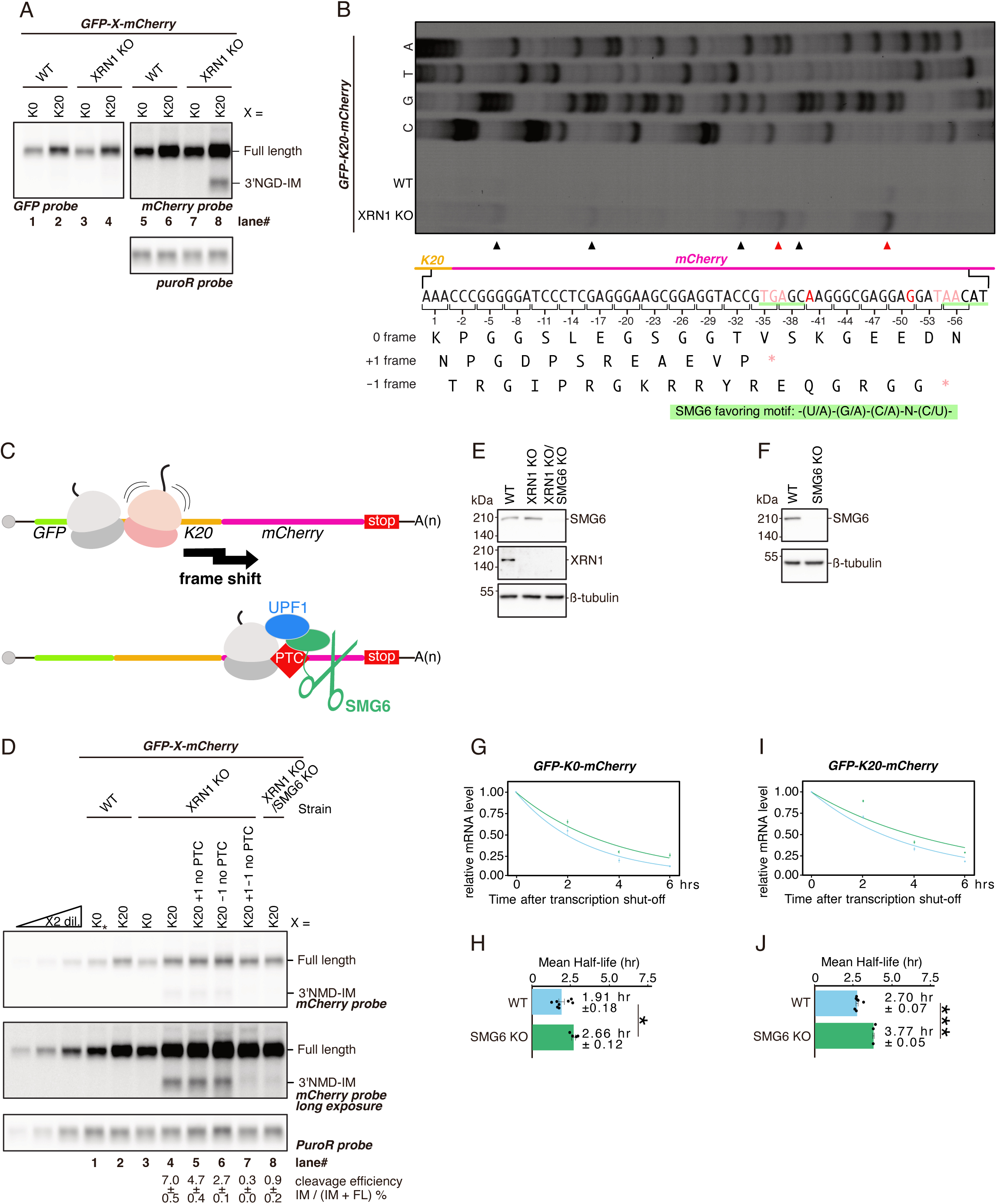
Ribosome slippage on K20 induces SMG6-dependent endonucleolytic cleavage of aberrant mRNAs. **A.** Northern blot analysis of *GFPX-mCherry* reporter mRNAs. mRNAs were detected using probes targeting the indicated regions: GFP for the 5′ end, mCherry for the 3′ end of the reporter. The *puroR* mRNA, expressed from an independent promoter, served as a transfection efficiency control. **B.** Primer extension analysis mapping the 5′ ends of the 3′ NGD intermediates. Red arrowheads indicate the major cleavage sites, corresponding to the nucleotides highlighted in vivid red. Pale red highlights premature termination codons (PTCs) arising in the +1 or -1 reading frames. An SMG6-favoring motif is indicated by a green underline. **C.** Proposed model for endonucleolytic cleavage induced by the K20 sequence. Translation of K20 sequence promotes ribosomal frameshifting, causing ribosomes to encounter a PTC and activate the nonsense-mediated decay (NMD) pathway, resulting in SMG6-dependent endonucleolytic cleavage of the aberrant mRNA. **D.** Northern blot analysis of the *GFP-K20-mCherry* reporter containing or lacking PTCs in the +1 and/or −1 reading frames. Asterisk indicates samples used for dilution. Cleavage efficiency was calculated as the ratio of the intermediate (IM) signal to the sum of intermediate and full-length (FL) signals. **E, F.** Western blot analysis of the cell lines used in this figure. Proteins were detected using antibodies against the indicated targets. **G, I.** Relative mRNA levels of the *GFP-X-mCherry* reporters normalized to the level of 0 h time point. Colours correspond to those used in H and J. **H, J.** Estimated mRNA half-lives calculated from the data shown in G and I. Each dot represents a half-life from individual experiment (n = 3–8); bars indicate mean values, and error bars represent S.E. Statistical analysis was performed using the Benjamini–Hochberg method. Adjusted *p* values are indicated as follows: *** *p*adj < 0.001; ** *p*adj < 0.01; * *p*adj < 0.05; N.S., not significant.

To gain insight into the mechanism underlying this cleavage, we next determined the precise cleavage sites by primer extension analysis and identified two major cleavage positions (Fig. 4B, red arrowheads). Unexpectedly, these cleavage sites were located downstream of the K20 collision sequence (highlighted in vivid red), in contrast to yeast NGD, in which the cleavage occurs within or upstream of collided ribosomes (Tomomatsu et al., 2023) (Fig.2F). Closer inspection of the sequence revealed that the cleavage sites were positioned in close proximity to premature termination codons (PTCs) present in either the +1 or -1 reading frame (highlighted in pale red). Given that poly-lysine sequences such as K20 are known to induce frequent ribosomal frameshifting (Chandrasekaran et al., 2019; Hashimoto et al., 2020; Juszkiewicz & Hegde, 2017; Koutmou et al., 2015; Tesina et al., 2020), we hypothesized that ribosome slippage on the K20 sequence causes ribosomes to enter alternative reading frames, encounter PTCs, and trigger SMG6-mediated endonucleolytic cleavage (Fig. 4C). Consistent with this model, an SMG6-favoring motifs were identified near the cleavage sites (underscored in green) (Schmidt et al., 2015) (Fig. 4B).

To test whether the cleavage depends on frameshift-induced PTC recognition, we constructed reporter variants lacking PTCs in the +1 frame, the -1 frame, or in both frames. 3′ NGD intermediates were still detected when a PTC was present in either frame alone but were completely absent when both frames lacked PTCs (Fig. 4D, lanes 4–7). Notably, cleavage efficiency was higher when only the -1 frame contained a PTC (Fig. 4D, lanes 5 and 6), consisting with both the primer extension results (Fig. 4B) and previously reported frameshifting biases that favour the -1 frame over the +1 frame in the context of K20 sequences (Arthur et al., 2015; Juszkiewicz & Hegde, 2017).

Based on these observations, we hypothesized that the endonucleolytic cleavage is mediated by SMG6 (Huntzinger et al., 2008), the endonuclease responsible for nonsense-mediated decay (NMD). To test this hypothesis, we examined cleavage in SMG6-deficient cells (Fig. 4E). Deletion of SMG6 completely abolished the 3′ NGD intermediate in XRN1/SMG6 double knockout cells (Fig. 4D, lane 8), demonstrating that cleavage is SMG6-dependent. The cleavage fragment was detected exclusively with the mCherry probe, confirming its identity as a 3′ cleavage product (Fig. S2B). Based on these results, we refer to this cleavage product as a 3′ NMD intermediate (3′ NMD-IM) (Fig. 4D). Finally, to assess the contribution of this endonucleolytic cleavage to overall mRNA degradation, we measured reporter mRNA half-lives using the Tet-off system in SMG6 knockout cells (Fig. 4F). Loss of SMG6 led to a modest stabilization of the K20 reporter (Fig. 4H–K), consistent with the relatively low efficiency of SMG6-mediated endonucleolytic cleavage observed by northern blot analysis (Fig. 4D).

## DISCUSSION

A major finding of this study is the first direct detection of a cleavage intermediate derived from a collision-inducing mRNA in mammalian cells (Fig.4B). Unexpectedly, this cleavage is mediated not by an NGD-specific nuclease but by SMG6 (Fig.4D), a key endonuclease of the nonsense-mediated decay (NMD) pathway (Huntzinger et al., 2008). As previously reported, ribosome slippage on the K(AAA)_20_ sequence induces frequent frameshifting (Arthur et al., 2015; Chandrasekaran et al., 2019; Hashimoto et al., 2020; Juszkiewicz & Hegde, 2017; Koutmou et al., 2015; Tesina et al., 2020), generating PTCs in alternative reading frames. These PTCs, in turn, trigger SMG6-dependent endonucleolytic cleavage, resulting in a distinct 3′ cleavage intermediate (Fig.4C).

While ribosome frameshifting on poly-A sequences has been extensively documented at the protein level (Arthur et al., 2015; Chandrasekaran et al., 2019; Hashimoto et al., 2020; Juszkiewicz & Hegde, 2017; Koutmou et al., 2015; Tesina et al., 2020), our study provides the first direct evidence linking such frameshifting events to mRNA cleavage and decay kinetics at the transcript level. These findings establish a mechanistic connection between ribosome slippage, NMD activation, and mRNA turnover at the transcript level. Importantly, ribosome frameshifting is not restricted to artificial reporters. Endogenous slippery sequences have been shown to promote programmed frameshifting in physiological contexts (Arthur et al., 2015). In addition, recent studies have shown that ribosomal frameshifting can occur on pseudouridine-modified mRNAs, including N^1^-methylpseudouridylated mRNA vaccines, leading to +1 frameshifting (Mulroney et al., 2024). Our results therefore suggest that SMG6-dependent cleavage triggered by ribosome slippage may represent a broader mechanism contributing to gene expression regulation beyond NGD reporters.

In this study, we further demonstrate that collision-dependent degradation of aberrant mRNAs is an evolutionarily conserved mechanism operating in both yeast and mammalian cells (Fig.1A). In both species, mRNAs undergoing ribosome stalling are degraded through two parallel pathways mediated by the ribosome collision sensor Hel2/ZNF598 and the Syh1–Smy2/GIGYF2 axis. Simultaneous disruption of both pathways resulted in pronounced stabilization of collision-inducing reporters, indicating functional redundancy (Fig.1C, 3D). Notably, in mammalian cells, the extent of stabilization observed in the ZNF598/GIGYF2 double knockout was less pronounced than that observed in yeast (Fig. 1C, 3D). This difference may reflect the multifunctional nature of GIGYF2 in mammals. GIGYF2 participates in multiple post-transcriptional regulatory pathways that are ultimately converged on conventional mRNA decay involving DCP2-mediated decapping and XRN1-dependent 5′–3′ exonucleolytic degradation (Zhao et al., 2025). In addition, GIGYF2 has been reported to repress nonsense-mediated decay (NMD) (Zinshteyn et al., 2021). In this study, we directly detected 3’NMD-IM resulting from frameshifting on the K20 sequence (Fig. 4D), which is mediated by the NMD endonuclease SMG6. Together, these findings raise the possibility of crosstalk between ribosome collision sensing and frameshift-induced NMD on slippery sequences.

Despite this conservation, we uncovered a major divergence between yeast and mammals in the role of endonuclease Cue2 and proposed endonuclease N4BP2. In yeast, Cue2-mediated cleavage is a central feature of NGD (D’Orazio et al., 2019; Müller et al., 2025; Tomomatsu et al., 2023), and our analysis further refined its contribution by distinguishing NGD–RQC⁺ and NGD–RQC⁻ cleavage modes (Fig.2F). We demonstrate that NGD–RQC⁻ cleavage plays a dominant role in mRNA decay, whereas NGD–RQC⁺ contributes only partially (Fig.2H). In contrast, we found no evidence that the proposed mammalian Cue2 homolog N4BP2 contributes to collision-dependent mRNA decay in our NGD reporter system. Although N4BP2 has been implicated in non-stop decay (NSD) contexts (Nicholson-Shaw et al., 2025), our results indicate that it is dispensable for NGD of collision-inducing mRNAs, at least under the conditions tested (Fig.3H).

This divergence raises several possibilities: Endonucleolytic cleavage during mammalian NGD may not be strictly required for mRNA decay, may be mediated by a distinct nuclease, or may be highly context dependent. Alternatively, the requirement for cleavage may depend on the nature and rigidity of collided ribosomes. Indeed, collisions induced by poly-lysine stretches (Chandrasekaran et al., 2019; Tesina et al., 2020) differs structurally from those induced by rare codons and *SDD1*-stalling sequence in yeast (D’Orazio et al., 2019; Ikeuchi et al., 2019; Matsuo et al., 2020), potentially influencing accessibility or affinity for endonucleases. Recent cryo-EM studies also have revealed that disome architectures vary depending on how ribosome collisions are induced (Huso et al., 2025), supporting the idea that distinct collision geometries may engage different downstream pathways.

We observed that the K20 reporter exhibited a slightly longer half-life than the non-stalling reporter in mammalian cells (Fig. 2D and F). One possible explanation is that ribosome-dense mRNAs are less accessible to mRNA decay machineries (Dilrangi et al., 2025) or are less efficiently routed to cytoplasmic decay sites such as P-bodies (Bicknell et al., 2024; El Fatimy et al., 2016; Ripin & Parker, 2022, 2023; Sheth & Parker, 2003). Consistent with this idea, we found that the K20 reporter mRNA was more enriched in polysome fractions (Fig. 3K and L).Moreover, live-cell imaging studies showing that ribosome stalling at poly(A) sequences promotes the formation of extended queues of collided ribosomes (Goldman et al., 2021) further support the model in which ribosome occupancy itself can protect mRNAs from degradation.

Finally, our findings have broader implications for viral infection. Both ZNF598 and GIGYF2 have been implicated in the control of viral mRNA translation (DiGiuseppe et al., 2018; J. Kim et al., 2025) as well as in innate immune responses (Wang et al., 2019). Given the widespread use of mRNA vaccines and the continuing threat of viral infectious diseases (J. Li et al., 2025; Nichol et al., 2000), a deeper understanding of translation-coupled mRNA quality control mechanisms may provide new avenues for therapeutic intervention and antiviral strategies.

## Methods

### Plasmid construction

Plasmids used in this study are listed in Table 1. DNA cloning and plasmid construction were performed using *Escherichia coli* DH5α. DNA fragments of interest were generated by PCR amplification by KOD-One (# KMM-101, TOYOBO) and inserted into linearized vectors by Gibson assembly or ligation using T4 DNA ligase (# M0202L, NEB). Oligonucleotides used for PCR were purchased from FASMAC. Primers used in this study are listed in Table 2. All cloned DNA fragments were verified by Sanger sequencing (Eurofins Genomics).

**Table 1.**
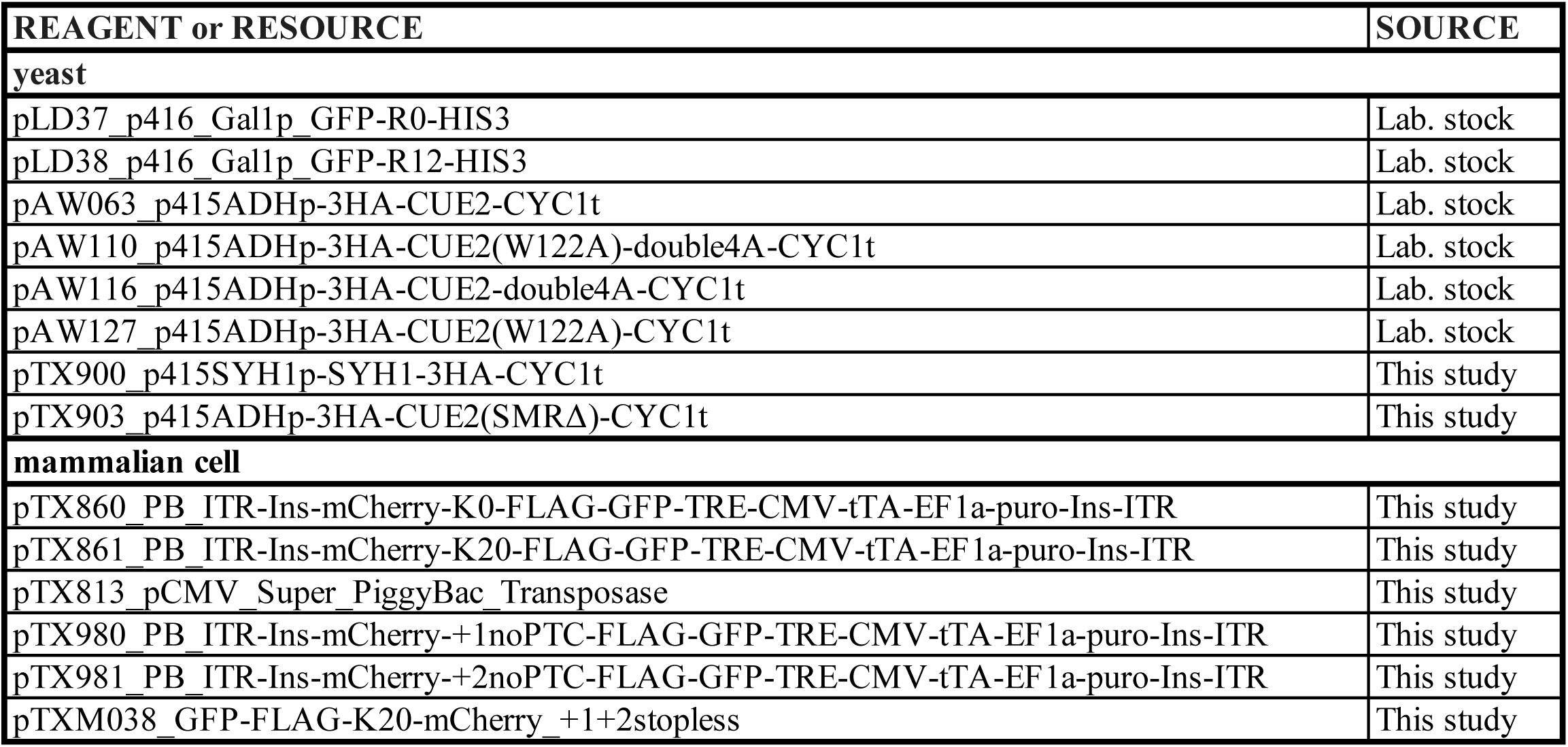
Plasmids used in this study.

**Table 2.**
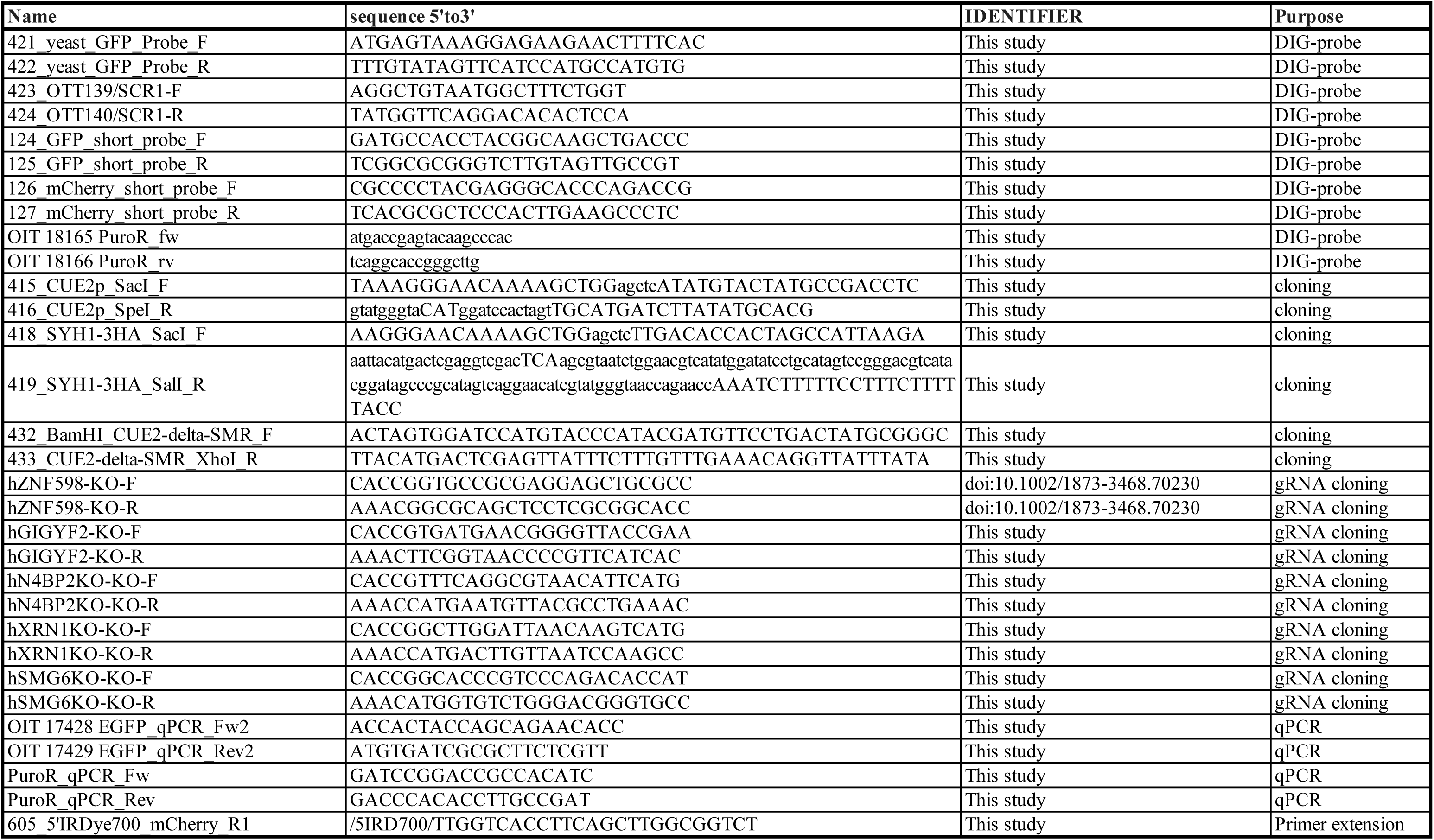
Primers used in this study.

The reporter plasmid *pTX860_PB_ITR-Ins-mCherry-K0-FLAG-GFP-TRE-CMV-tTA-EF1α-puro-Ins-ITR* was constructed using the PB-CMV-MCS-EF1-Puro backbone (#PB510-B-1, System Biosciences). The tTA fragment was amplified from cDNA prepared from the HEK293 Tet-Off® Advanced Cell Line (#631152, Clontech) and inserted into the MCS in PB-CMV-MCS-EF1-Puro. The TRE cassette derived from the pTet-Off Advanced vector (#631070, Clontech) and the *GFP–FLAG–K0–mCherry* reporter cassettes from *pmGFP-P2A-K0-P2A-RFP*, a gift from Dr. Ramanujan Hegde (Addgene plasmids #105686 (Juszkiewicz & Hegde, 2017)) were inserted into SpeI site between Core Insulator and CMV promoter. *pTX813_pCMV_Super_PiggyBac_Transposase* was constructed based on the Super PiggyBac Transposase Expression Vector (#PB210PA-1, System Biosciences).

### Yeast strains and culture

*Saccharomyces cerevisiae* strains used in this study were W303-1a (ATCC stock number, 208352; genotype: MATa ade2-1 ura3-1 his3-11 trp1-1 leu2-3 leu2-112 can1-100) and its derivatives, listed in Table 3. Gene disruption strains and genomic tagged strains were constructed by established homologous recombination strategies using polymerase chain reaction (PCR)-amplified selection marker genes *KanMX4*, *HphMX4*, *NatMX4*, *3HA-HphMX4*, *3HA-His3MX6*, or *His6-TEV-ProteinA (HTP)*-*His3MX4 (Janke et al., 2004; Longtine et al., 1998)*. Cells with the gene of interest deleted or tagged were selected on plates containing G-418 (# 074-05963, Wako), Hygromycin B (# 085-06153, Wako), or Nourseothricin (# AB-101S, JenaBioScience), or lacking histidine.

**Table 3.** Yeast strains used in this study.

| Strain# | Genotype/helper plasmid | Source |
| --- | --- | --- |
| W303-1a | *MATa leu2-3,112 trp1-1 can1-100 ura3-1 ade2-1 his3-11,15* | Lab stock |
| YKI110 | *W303-1a, hel2Δ::natMX4* | https://doi.org/10.1038/s41467-017-00188-1 |
| YYM113 | *cue2Δ::kanMX4* | https://doi.org/10.1093/nar/gkac1172 |
| ST146 | *W303 syh1Δ::natMX4 smy2Δ::kanMX4* | This study |
| ST147 | *W303 syh1Δ::natMX4 hel2Δ::kanMX4 smy2Δ::hphMX6* | This study |
| YJZ004 | *W303 syh1D::natMX4 smy2D::kanMX4 cue2D::hygMX6* | This study |

In general, yeast cells were grown in yeast extract peptone (YP) or synthetic complete (SC) medium with 2% glucose. Cells carrying *GFP-R0/R12-HIS3* plasmids were grown in SC medium containing 2% galactose to induce transcription from the *GAL1* promoter. Cells were incubated with shaking at 30 °C to the log phase (optical density at 600 nm; OD600 = 0.5 to 0.8), harvested by centrifugation at room temperature, and flash frozen in liquid nitrogen unless otherwise noted.

To evaluate the stability of aberrant mRNA derived from the *GAL1p- GFP-R0/R12-HIS3* plasmids, cells were grown in SC medium containing 2% galactose to an OD600 of 0.5. A 15 mL culture was centrifuged at 3500 × *g* for 2 min, and the cell pellet was re-suspended in 15 mL of pre-warmed SC medium containing 2% glucose to shut off transcription from the *GAL1* promoter. The re-suspended culture was continuously incubated, and 1.5 mL was harvested at various timepoints for northern blot analysis. Cells harvested immediately after transcription shut-off were designated as the 0 min samples(S. Li et al., 2025).

### Sucrose density gradient centrifugation-Western Blotting (SDG-WB) for yeast

Cells were ground in liquid nitrogen and suspended on ice in SDG lysis buffer (20 mM HEPES-KOH pH 7.4, 100 mM KOAc, 2 mM Mg(OAc)_2_, 100 μg/mL CHX, 1 mM dithiothreitol (DTT), 1 mM phenylmethanesulfonyl fluoride (PMSF), cOmplete Mini EDTA-free Protease Inhibitor Cocktail (#11836170001, Roche) 1 tablet/10 mL) at a volume proportional to (OD600/0.6) × culture volume (mL) × 5–10 µL. After centrifugation at 20,000 × g for 30 min at 4 °C, the clear supernatants were collected as cell lysates. Lysates (total RNA amount equivalent to 200µg) were layered onto 10%–50% sucrose gradients in 10 mM Tris-acetate pH 7.5, 70 mM NH_4_OAc, and 4 mM Mg(OAc)2, prepared in 14 × 95 mm Seton PolyClear™ tubes (# 151-514B) using a Gradient Master (Biocomp). Centrifugation was performed at 283,807 × g (40,000 rpm) in a P40ST rotor (HITACHI) for 2 hr at 4 °C. Fractions were collected from the top of the gradient using a Piston Gradient Fractionator™ (BioComp) while continuously monitoring absorbance at 260 nm with a single path UV-1 optical unit (ATTO Biomini UV-monitor) connected to a chart recorder (ATTO digital mini-recorder). Each of the collected fractions were mixed with TCA (#34605-95, Nakarai, final 10%v/v), and centrifuged at 20,000 × g for 30 min at 4 °C. The pellet was washed with acetone (#00309-35, Nakarai) and resuspended in 50uL of 4XSB not pHed (3.5% SDS, 14% glycerol, 120mM Tris, 8mM EDTA, 120mM DTT, 0.01% Bromophenol Blue).

Samples were heated at 95 °C for 5 min. Proteins were separated on 12% SDS–polyacrylamide gels and transferred onto PVDF membranes (Immobilon-P, Millipore) using a semi-dry transfer system. Membranes were blocked with 5% skim milk and incubated with antibodies. After washing, membranes were treated with ImmunoStar LD (# 290-69904, Wako), and chemiluminescence was detected by ImageQuant LAS4000 mini (GE Healthcare). The following primary antibodies were used: Anti-HA-Peroxidase (# 12013819001, Roche, RRID: AB_390917), 1/3000.

### RNA extraction and northern blot for yeast

Total RNA was isolated from frozen cells using the hot phenol RNA extraction method as follows: Cells were re-suspended with 200 μL of RNA buffer (Tris-HCl pH 7.5, 300 mM NaCl, 10 mM EDTA, 1% SDS) on ice and mixed with 200 μL of water-saturated phenol. The mixture was incubated at 65 °C for 5 min, vortexed for 10 sec, and then chilled on ice for 5 min. Following centrifugation at 16,000 × *g* for 5 min at room temperature, 200 μL of the aqueous phase was collected and mixed with 200 μL of water-saturated phenol/chloroform/isoamylalcohol (25:24:1). After a second centrifugation at 16,000 × *g* for 5 min at room temperature, 180 μL of the aqueous phase was collected, combined with 18 μL of 3 M NaOAc pH 5.2 and 450 µL of ice-cold ethanol, and incubated at − 80 °C for 30 min. RNA was precipitated by centrifugation at 20,000 × *g* for 15 min at 4 °C, and re-suspended in 20 µL of RNase free water.

For RNA sample preparation, 6 µL of RNA solution was mixed with 12.5 μL of deionized formamide, 2.5 μL of 10 x MOPS buffer pH 7.0 (0.2 M MOPS, 10 mM EDTA, 50 mM NaOAc), 4 μL of 37% formaldehyde, and 2.5 µL of RNA dye (50% glycerol, 10 mM EDTA pH 8.0, 0.05% bromophenol blue, 0.05% xylene cyanol). RNA samples were heated at 65 °C for 5 min, then chilled on ice for 5 min. Samples were separated on 1.2% agarose-formaldehyde gels, transferred to Hybond-N+ membranes (GE healthcare) using a capillary system, and cross-linked to membranes using a CL-1000 ultraviolet crosslinker (UVP) at 120 mJ/cm^2^. Membranes were pre-incubated with DIG Easy Hyb Granules (# 11796895001, Roche) and hybridized with DIG-labeled GFP and SCR probes. DIG-labelled probes were prepared using PCR DIG Probe Synthesis Kit (#11636090910, Roche). Primers used for generating probes are listed in Table 2. After hybridization, membranes were washed, blocked with Blocking Reagent (# 11096176001, Roche), and incubated with Anti-Digoxigenin-AP, Fab fragments (# 11093274910, Roche). Membranes were then washed, equilibrated to pH 9.5, and treated with CDP-star (# 11759051001, Roche). Chemiluminescence was detected using ImageQuant LAS4000 mini. Band intensities were quantified using ImageQuant TL software (Cytiva).

### Cell strain and culture

HEK293T (CVCL_0063) cells were provided by the RIKEN BRC through the National Bio-Resource Project of MEXT, Japan, and cultured in low glucose Dulbecco’s modified Eagle’s medium, DMEM (#08456-36; Nacalai Tesque), supplemented with 10% fetal bovine serum (FBS, #556-33865, Biosera) and 100 U / mL penicillin/streptomycin (PS, # 09367-34; Nacalai Tesque). HEK293T cells were purchased from the RIKEN BRC and Thermo Fisher individually in the past 3 years and were not authenticated. All cells were regularly confirmed to be free of myco-plasma contamination using DAPI staining (Huang et al., 2025).

### Construction of KO cells

Plasmids containing guide RNAs were transfected into cells, and single-cell clones were obtained by limiting dilution. The resulting clones were genotyped by Western blotting, and genomic edits were analysed by Sanger sequencing. The gRNA sequence was designed using CHOPCHOP v3 (Labun et al., 2019) and inDelphi (Shen et al., 2018), genome sequence was analysed by CRISP-ID (Dehairs et al., 2016). pSpCas9(BB)-2A-Puro (PX459), a gift from Feng Zhang (Addgene plasmid #62988 (Ran et al., 2013)) or SpCas9-HF1-plus, a gift from Feng Welker E (Addgene plasmid #126768 (Kulcsár et al., 2020)) were used for genome editing.

### Plasmid transfection

When HEK293T cells had grown to about 80 % confluence in a 12-well plate, we prepared a transfection mixture by combining PEI-MAX (#24765-1, Cosmo Bio,) and plasmid DNA at a 3 : 1 ratio in Opti-MEM™ Reduced-Serum Medium (#31985-062, Thermo Fisher). 1μg of DNA was used for transfection. The mixture was then added dropwise to the cells. After a 3 h incubation at 37 °C in a 5 % CO₂ atmosphere, the transfection medium was replaced with fresh DMEM.

### Preparation of cell lysates for WB and WB for cell

Cells were washed once with PBS and lysed in RIPA buffer (20 mM Tris-HCl [pH 7.5], 150 mM NaCl, 2 mM EDTA, 10 mM sodium fluoride, 10 mM β-glycerophosphate, 1 % Triton X-100, 0.1 % SDS, 0.5 % sodium deoxycholate, 40 mM N-ethylmaleimide). Protein concentrations were determined with the Protein Assay Dye Reagent Concentrate (#5000006, Bio-Rad).

Lysates were mixed with distilled water and 4XSB (375 mM Tris–HCl [pH 6.8], 50% glycerol, 9% SDS, 0.03% Bromophenol Blue, 6% 2-Mercaptoethanol) to the final protein concentration of 2μg/μL. Then samples were boiled at 95 °C for 5min to denature the proteins. WB was performed as described in the “SDG-WB for yeast” section. The following primary and secondary antibodies were used:

Anti-ZNF598 (# A305-108A, Thermo Fisher Scientific, RRID: AB_2631503), 1/3000 dilution;

Anti-GIGYF2 (# 24790-1-AP, Proteintech, RRID: AB_2879727), 1/3000 dilution; Anti-SMG6 (# ab87539, Abcam: AB_10674461), 1/3000 dilution;

Anti-XRN1 (# sc-165985, Santa Cruz Biotechnology: AB_2304774), 1/3000 dilution; Anti-β-tubulin (# 014-25041, FUJIFILM Wako Pure Chemical Corporation: AB_3717508), 1/3000 dilution;

Anti-N4BP2 (# 23892-1-AP, Proteintech: AB_3669422), 1/3000 dilution; ECL Anti-Rabbit IgG (# NA934, GE Healthcare, RRID:AB_772206); and ECL Anti-Mouse IgG (# NA931, GE Healthcare, RRID: AB_772210) conjugated with horseradish Peroxidase (HRP), 1/5000 dilution.

### RNA extraction and northern blotting for cell

Total RNA from HEK293T cells was extracted using ISOGEN II according to the manufacturer’s instructions (#311-07361, NIPPON GENE). For primer extension analysis, RNA was treated with DNase I (#AMPD1-1KT, Sigma-Aldrich) and purified by phenol/chloroform/isoamyl alcohol (25:24:1) method as described in “RNA extraction and northern blotting for yeast” section. A total of 3 µg of RNA was used for northern blot analysis. Subsequent procedures were performed as described in the “RNA extraction and northern blotting for yeast” section.

DIG-labeled probes used for detection are GFP short, mCherry short, and puroR probes. Primers used for probe generation are listed in Table 2.

### Half-life measurement by Tet-off system

For half-life measurements, HEK293 cells stably expressing reporter were generated using the PiggyBac Transposon Vector System (Ding et al., 2005; A. Kim & Pyykko, 2011)(System Biosciences). HEK293T cells in 6-cm dishes were cultured at approximately 80% confluence and transfected with reporter plasmid, transposase plasmid, and PEI-Max at a ratio of 3 μg : 1.2 μg : 4.2 μL, respectively. Three hours after transfection, the medium was replaced with fresh medium containing 1 μg/mL doxycycline (#049-31121, FUJIFILM Wako Pure Chemical Corporation), and cells were incubated for 24 h. Cells were then selected with 1 μg/mL puromycin (#166-23153, FUJIFILM Wako Pure Chemical Corporation) for 24 h. Subsequently, cells were maintained in medium containing 1 μg/mL doxycycline for 5 days to eliminate cells retaining episomal plasmids. Cells were then trypsinized and washed with DMEM without doxycycline, seeded into 24-well plates at a density of 5.0 × 10⁴ cells per well, and cultured for 24 h. The medium was then replaced with pre-warmed medium containing 1 μg/mL doxycycline, and cells were harvested at the indicated time points.

Total RNA was extracted using ISOGEN II according to the manufacturer’s instructions (#311-07361, NIPPON GENE). A total of 125–500 ng RNA was used for reverse transcription and quantitative PCR (qPCR) analysis. RT-qPCR was performed with RNA-direct® Realtime PCR Master Mix (#QRT-101, TOYOBO). Reverse transcription (RT) and qPCR were performed using the following thermal cycling conditions: Reverse transcription: denaturation at 90°C for 30 s; reverse transcription at 60°C for 20 min; denaturation at 95°C for 1 min. PCR amplification: denaturation at 95°C for 15 s, annealing at 51°C for 15 s, and extension at 74°C for 45 s, for 45 cycles. RT-qPCR was performed using the CFX Connect Real-Time System (Bio-Rad). Primer sequences used for qPCR are listed in Table 2.

### Sucrose density gradient centrifugation-Northern Blotting (SDG-NB)

HEK293T cells cultured in 10-cm dishes were transfected at approximately 80% confluence with 24 μg of reporter plasmid and incubated for 24 h. Cells were collected by scraping and washed once with PBS. After the removal of PBS, cells were lysed by 500 μL of lysis buffer (50 mM Tris-HCl, pH 6.8; 100 mM NaCl; 10 mM MgCl₂; 1% NP-40; 100 μg/mL cycloheximide; 2 mM 2-mercaptoethanol; 1 mM phenylmethanesulfonyl fluoride; cOmplete Mini EDTA-free Protease Inhibitor Cocktail [#11836170001, Roche], 1 tablet per 10 mL) with gentle pipetting and incubated on ice for 5 min. Cell lysates were clarified by centrifugation at 1,500 × g for 5 min at 4°C, and the supernatant was collected. RNA concentration was measured using a NanoDrop spectrophotometer (Thermo Fisher Scientific). Lysate corresponding to 200 μg of total RNA was layered onto 10–50% sucrose gradients in 20 mM HEPES-NaOH, pH 6.8; 100 mM NaCl; 10 mM MgCl₂ prepared in 14 × 95 mm Seton PolyClear™ tubes (#151-514B) and a Gradient Master (Biocomp). Gradients were centrifuged at 283,807 × g (40,000 rpm) in a P40ST rotor (HITACHI) for 2 h at 4°C.

RNA was isolated from sucrose gradient fractions using a guanidine-HCl precipitation method. Immediately after fractionation, 226 μL of each fraction was mixed with 500 μL of 8 M guanidine-HCl and 750 μL of ethanol, followed by incubation at −30°C overnight. RNA was precipitated by centrifugation at 20,000 × g for 20 min at 4°C, washed with ice-cold 75% ethanol, and resuspended in 200 μL RNA buffer. Samples were then mixed with 20 μL of 3 M sodium acetate (pH 5.2) and 600 μL ice-cold ethanol, incubated at −30°C for 1 h, and centrifuged at 20,000 × g for 20 min at 4°C. RNA pellets were washed with ice-cold 75% ethanol and resuspended in 10 μL RNase-free water. The recovered RNA was subsequently used for Northern blotting, as described in the section “RNA extraction and Northern blotting for cells”.

### Half-life calculation and statistical analysis

Quantified mRNA level of GFP was normalized to internal controls: SCR RNA in yeast and puroR mRNA in mammalian cells. RNA abundance at each time point was normalized to the initial (0 h) level. mRNA decay kinetics were fitted using a single-exponential decay model:

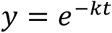

where *k* represents the decay constant, t represents the time after transcription shut off, y represents the normalized RNA level. Curve fitting was performed using nonlinear least-squares regression implemented in R (nls function). The mRNA half-life (*t*₁/₂) was calculated according to the equation derived from a first-order decay equation (Chen et al., 2008):

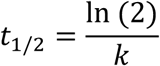

Outliers were identified and excluded using the interquartile range (IQR) method. Values lying more than 1.5 × IQR above the third quartile or below the first quartile were considered outliers and removed prior to statistical analysis.

### Primer extension analysis

Total RNA (240 µg) was subjected to reverse transcription using SuperScript IV Reverse Transcriptase (Invitrogen, Cat# 18090010) with a 5′-IRDye700–labeled primer (IDT) complementary to the mCherry sequence. An equal volume of phenol/chloroform/isoamyl alcohol (25:24:1) was added to the reaction mixture, followed by vortexing and centrifugation. The aqueous phase was collected, and cDNA was precipitated by adding two-fifths volume of 5 M NH₄OAc, an equal volume of isopropanol, and 1 µL of glycogen. The resulting cDNA pellet was dissolved in 8 µL of stop solution without xylene cyanol (95% formamide, 20 mM EDTA pH 8.0, 0.01% bromophenol blue). Samples were denatured at 70 °C for 2 min, immediately chilled on ice for 5 min, and separated on a 5% polyacrylamide–TBE–urea sequencing gel by electrophoresis at 1000 V for 120 min. IRDye700 fluorescence was detected using an Amersham Typhoon scanner 5 system (Cytiva). The sizes of reverse transcription products were determined by comparison with a sequencing ladder generated from the corresponding reporter plasmid DNA using the same primer and the Thermo Sequenase Cycle Sequencing Kit (# 78500 1KT, USB).

### Statistics information

All details on quantification and statistical analysis of experiments can be found in the figure legends. All statistical tests were performed using R 4.2.2 (https://www.r-project.org/), Rstuido 2025.9.2.418 (http://www.posit.co/). Further details are provided above for each type of experiment in the corresponding sections.

### Reporting summary

Further information on research design is available in the Nature Portfolio Reporting Summary linked to this article.

## Acknowledgements

We would like to thank Dr. Sihan Li, Dr. Toru Suzuki, Mr. Hideaki Takeda for helpful discussions and critical reading of the manuscript. We thank Miho Hoshi for construction of plasmids. This work was supported by the following funding: This work was supported by This work was supported by AMED (JP23gm1110010, JP223fa627001 to TI), MEXT/JSPS KAKENHI (Grant Numbers 22H0040, 25H000071 to T.I., 25K18399 to S.H.), Research grants from Takeda Science Foundation (T.I.) and Mitsubishi Foundation (T.I.).

## Author contributions

S.H. and T.I. conceptualized the study. S.H. developed the methodology and carried out data visualization. S.H. and J.S. performed the investigation. J.Z. and C.K. provided essential resources. S.H. wrote the original draft of the manuscript. S.H. and T.I. reviewed and edited the manuscript. T.I. supervised the project and managed project administration.

## Additional information

List of materials used in this study.

**Correspondence and requests for materials should be addressed to**

Satoshi Hashimoto or Toshifumi Inada.

---

**Fig. S1. Northern blot images used for mRNA half-life calculations**

The reporter mRNAs were detected using a GFP probe. SCR RNA was used as a loading control. Red boxes indicate the northern blot images presented in the main figures.

**Fig. S2. Mapping of endonucleolytic cleavage sites**

**A.** Full-length gel image corresponding to the primer extension analysis shown in Fig. 4B. The red box indicates the region presented in the main figure.

**B.** Northern blot shown in Fig. 4F probed with GFP. The same samples were analyzed. No cleavage intermediates were detected on the 5′ side of the reporter mRNA.

