## Supplementary figures and images for "Parallel ZNF598 and GIGYF2 pathways mediate conserved collision-dependent mRNA decay with distinct endonucleolytic features"

### Supplemental Figures

Hashimoto *et al.*, Fig. S1

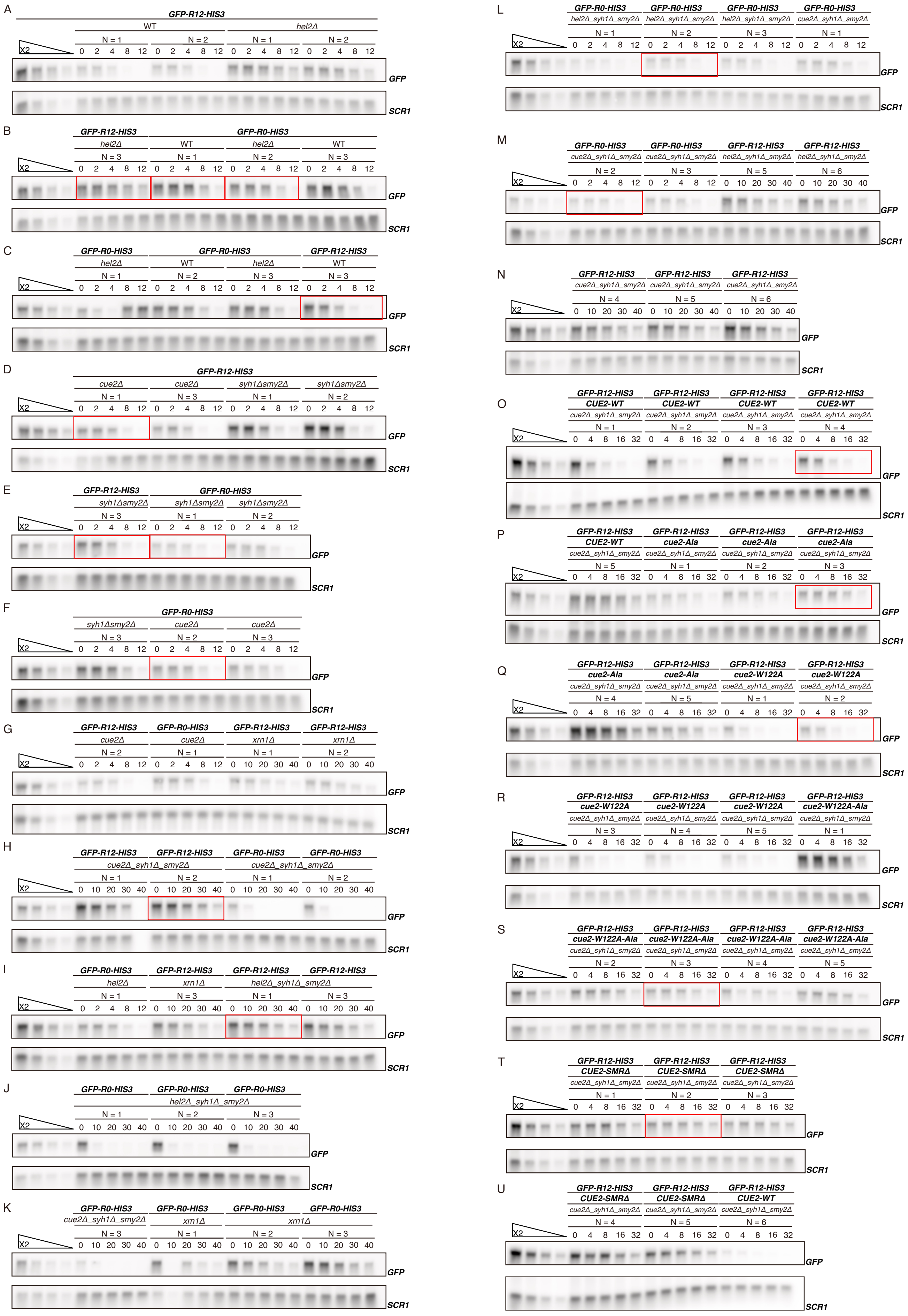

Hashimoto *et al.*, Fig. S2

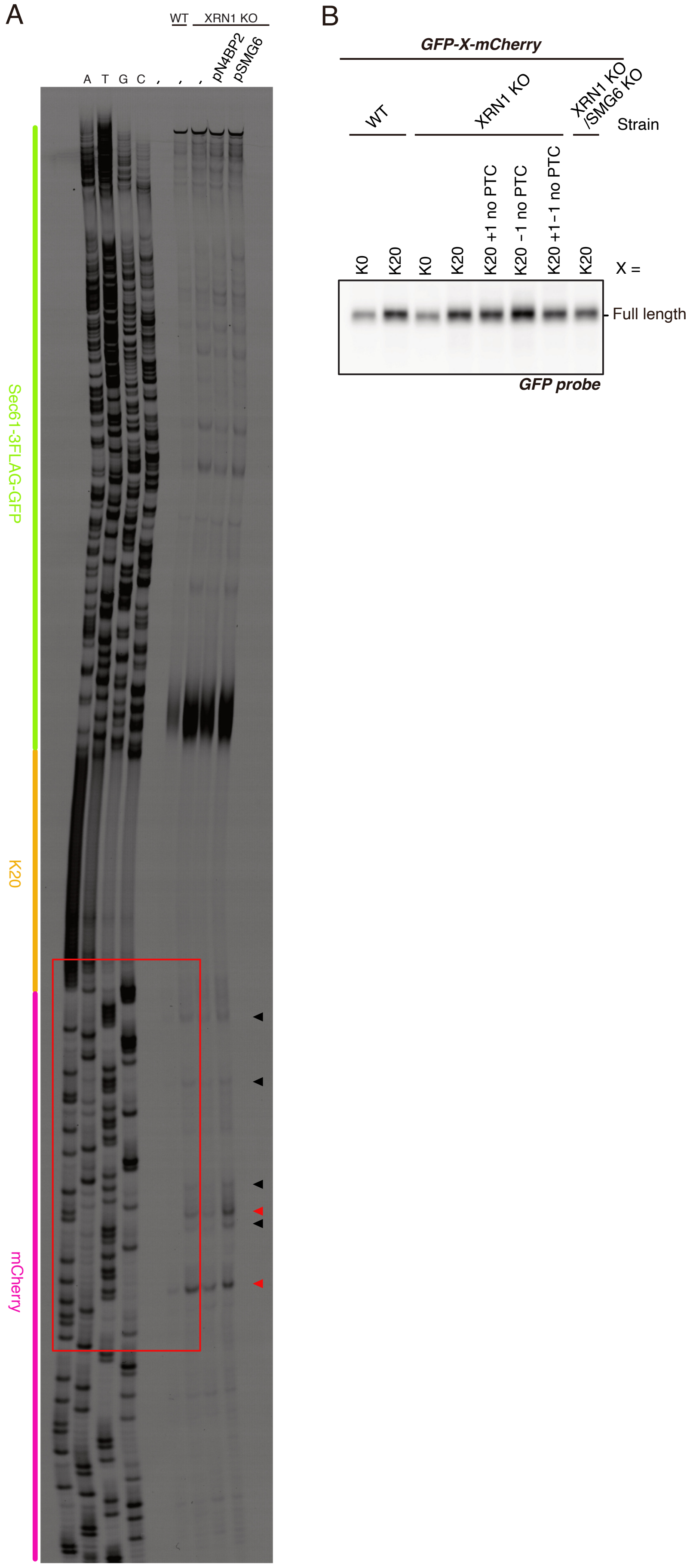
